# Comparative transcriptomics reveals differences in cortical cell type organization between metatherian and eutherian mammals

**DOI:** 10.1101/2025.10.03.680397

**Authors:** Ryan Gorzek, Joshua T. Trachtenberg

## Abstract

The neocortex, a layered structure unique to mammals, supports higher-order functions including perception, learning, and decision-making. While its laminar architecture is broadly conserved, the cell type-specific organization of the cortical column has not been compared across species that diverged early in mammalian evolution. To address this, we used single-nucleus RNA sequencing and spatial transcriptomics to compare gene expression, cell types, and laminar architecture in the primary visual cortex (V1) of metatherian (*Monodelphis domestica*) and eutherian (*Mus musculus*) mammals. We show that spatio-transcriptomic distinctions between supragranular (layer 2/3) and infragranular (layer 5) intratelencephalic (IT) neurons are more pronounced in mice, consistent with lineage-specific specialization. Mouse cortex also exhibits a lower relative density of parvalbumin-positive (PV) GABAergic neurons and redistributed perineuronal nets, consistent with altered constraints on plasticity. Together, these findings demonstrate substantial variation in the cellular and spatial organization of the cortical column across deeply diverged mammals, challenging the view that local cortical architecture is uniformly conserved.

**Significance Statement:** The neocortex supports perception and flexible behavior in mammals, yet how its cellular composition varies across early mammalian divergences has not been directly examined. By comparing transcriptomic cell types and their spatial organization in the primary visual cortex of a metatherian (opossum) and a eutherian (mouse), we show that major classes and laminar structure are broadly conserved, but intratelencephalic neurons differ substantially. These differences are accompanied by shifts in inhibitory circuitry and extracellular scaffolding, indicating divergent modes of cortical organization and plasticity across mammalian lineages. Our findings challenge the assumption that neocortical columns are uniformly conserved and identify specific cellular populations that vary across deeply diverged therian mammals.

## Introduction

The neocortex is a hallmark of mammalian evolution, enhancing perception and action through its characteristic laminar and columnar organization (*1, 2*). Central to its evolutionary elaboration is cortical arealization, the subdivision of cortex into specialized functional regions that scale computational capacity as brains enlarge (*3–6*). By contrast, comparative studies of cellular organization along the cortical depth—often termed a column—have emphasized the conservation of laminar structure, projection patterns, and major neuronal subclasses (*7, 8*). This has shaped the prevailing view that columnar architecture is conserved across mammalian lineages (*9*), with primate V1 representing a notable exception (*10–12*). Whether core features of the cortical column vary across the earliest divergences in mammalian evolution has not been directly examined.

The earliest divergence within Theria—the split between metatherians (marsupials) and eutherians (placentals) ∼160–180 million years ago (*13–15*)—provides an opportunity to examine differences in cortical organization across a deep phylogenetic divide. The gray short-tailed opossum (*Monodelphis domestica*), a metatherian mammal, retains many features often interpreted as reflecting the ancestral therian brain, including prolonged postnatal development, small and unlaminated thalamic nuclei, and a lissencephalic neocortex (*16–21*). However, like all extant mammals, opossums represent the outcome of tens of millions of years of independent evolution and cannot be treated as proxies for ancestral states.

Because the neocortex evolves under lineage-specific developmental and ecological constraints, comparisons between extant mammals cannot reconstruct ancestral cortical organization or localize changes to a specific divergence point. Instead, such comparisons can identify conserved and labile features, constraining hypotheses about evolutionary stability in cortical organization. Comparative transcriptomic studies have established relationships among neocortical cell types within Euarchontoglires (rodents and primates) (*22*) and between the neocortex and the dorsal pallium of birds, reptiles, and amphibians (*23–27*). However, cortical cell type organization in metatherian mammals remains largely unexplored, limiting our understanding of the conservation of neocortical cell types and architecture across Theria.

To address this gap, we used single-nucleus RNA sequencing (snRNA-seq) and spatial transcriptomics (Stereo-seq) to compare the primary visual cortex (V1) of *M. domestica* and the laboratory mouse (*Mus musculus*). Mice were selected as a placental comparator based on both biological and practical considerations; in contrast to primates, mice share several broad life-history and ecological features with opossums, including nocturnality, short lifespan, small body size, and reliance on sensory-guided foraging under predation pressure. These shared constraints reduce the likelihood that differences in cortical organization arise solely from disparities in sensory ecology or longevity. Accordingly, mice serve as a well-characterized reference point within Eutheria, rather than a proxy for placental mammals as a whole. Retinal cell types have been characterized in both species, revealing broadly conserved classes of retinal ganglion cells and thus comparable visual information streams relayed through the thalamus to V1 (*28*).

Although observed differences may reflect lineage-specific adaptations and do not support clade-level generalization, our data show that intratelencephalic (IT) glutamatergic neurons differ markedly in transcriptomic identity and spatial organization between opossums and mice, despite broad conservation of major neuronal subclasses and laminar organization. Consistent with the concept of evolutionary specialization and division of labor (*29–31*), opossum V1 exhibits more generalized IT neuron identities, whereas mouse V1 displays sharper spatio-transcriptomic distinctions among IT populations. In addition, differences in inhibitory interneuron populations and perineuronal net distribution suggest altered constraints on cortical plasticity, including shifts in magnitude and laminar localization (*32–34*). Together, these findings demonstrate that substantial variation in cortical cellular architecture exists across deeply diverged mammalian lineages and highlight IT neuron populations as a major axis of evolutionary divergence.

## Results

### A single-nucleus transcriptomic atlas of the primary visual cortex in the gray short-tailed opossum

We generated a single-nucleus RNA sequencing (snRNA-seq) atlas of the adult gray short-tailed opossum (*Monodelphis domestica*) V1 using 10x Chromium (Fig. 1A, B, see Methods). Tissue from both hemispheres was collected from two animals, pooled by animal, and used to generate four transcriptomic libraries. After preprocessing (see Methods, fig. S1H–S), we obtained 32,764 high-quality nuclear transcriptomes (Fig. 1C). We compared these data to our previously reported snRNA-seq dataset from adult (P38) mice (*35*), generated using similar protocols (Fig. 1D).

**Figure 1.**
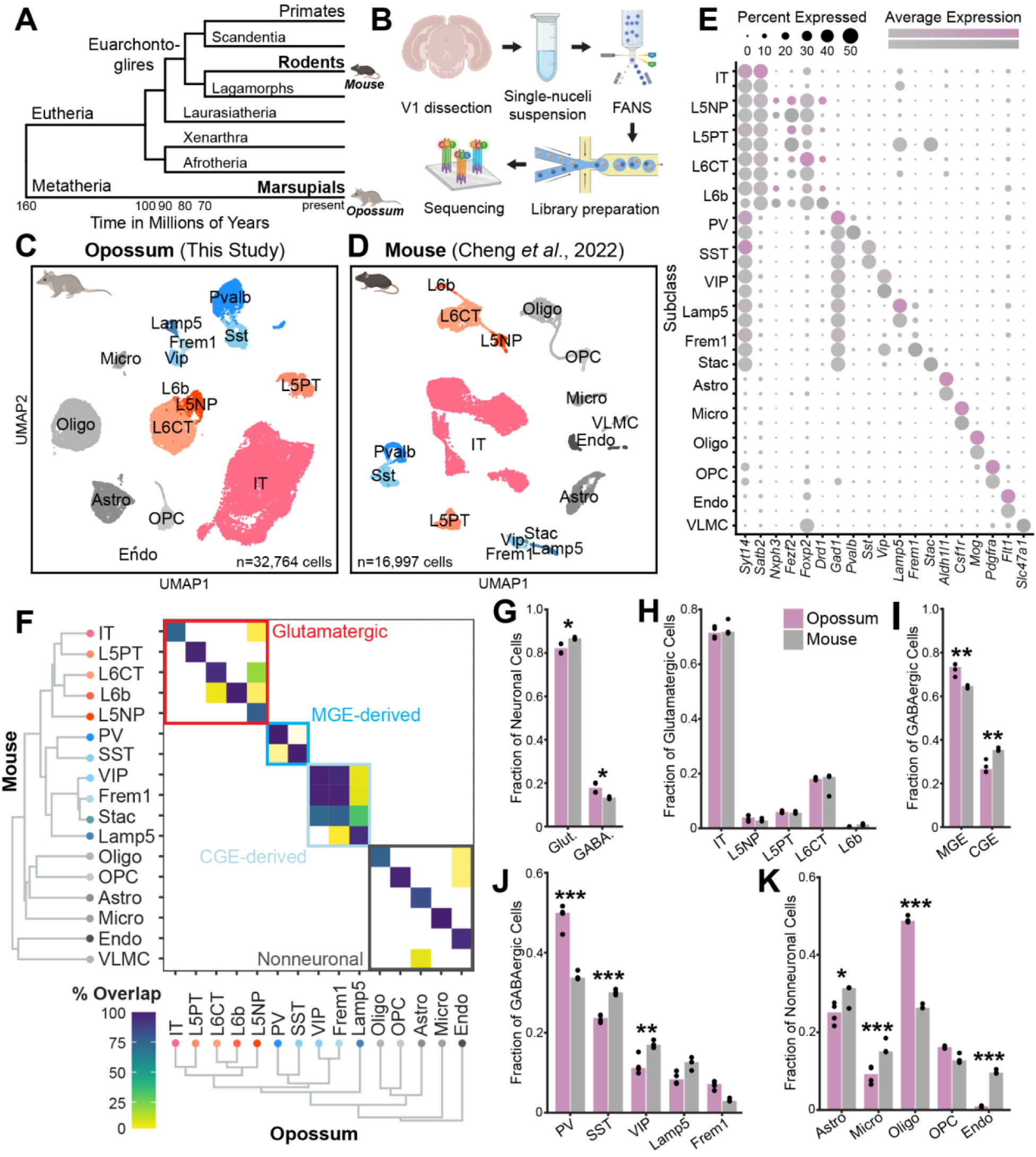
Cross-species transcriptomic comparison in primary visual cortex (V1) of the metatherian gray short-tailed opossum (*Monodelphis domestica*) and eutherian mouse. (**A**) Cladogram of therian mammalian evolution. (**B**) Experimental workflow of single-nucleus RNA sequencing (snRNA-seq) in opossum V1. (**C**) Opossum cells embedded in uniform manifold approximation (UMAP) dimensions, labeled by subclass. (**D**) Same as (C) but mouse cells from our previous study (*35*). (**E**) Canonical marker gene expression used in part to identify opossum subclasses. (**F**) Mouse and opossum subclass-level dendrograms and co-clustering in an integrated cross-species transcriptomic space. (**G–K**) Relative proportions of some transcriptomic groupings (G, class; H, glutamatergic subclass; I, GABAergic origin structure; J, GABAergic subclass; K, non-neuronal subclass) are significantly different between species (ANOVA followed by two-sided Tukey’s HSD test and Bonferroni correction; *p < 0.05, **p < 0.01, ***p < 0.001; table S1). Opossum data points include two biological replicates with two technical replicates each, mouse data points include three biological replicates.

Following dimensionality reduction and clustering, we used canonical marker genes (Fig. 1E, fig. S1L–M) and whole-transcriptome classification (fig. S1Q–S) to define major cell classes—glutamatergic, GABAergic, and non-neuronal—and their subclasses in opossum V1. Cross-species integration and co-clustering revealed that most subclasses were conserved, and transcriptomic hierarchies were broadly similar across species, consistent with conserved cellular taxonomies (Fig. 1F). However, subclass-level frequencies diverged. While glutamatergic neurons were more abundant in the mouse (Fig. 1G), the relative proportions of glutamatergic subclasses were nearly identical (Fig. 1H). By contrast, GABAergic composition differed markedly: opossum inhibition was biased towards medial ganglionic eminence (MGE)-derived interneurons that express *Pvalb* or *Sst* (Fig. 1I), with parvalbumin-positive (PV) cells comprising nearly 50% of the GABAergic population—far more than in mouse (∼35%, Fig. 1J; confirmed by histology, Fig. 4A–C). In mice, inhibition was biased towards caudal ganglionic eminence (CGE)-derived subtypes that express *Vip*, *Lamp5*, or *Frem1* (Fig. 1I). Notably, MGE-derived interneurons typically participate in feed-forward inhibition, driven by early thalamic and intracortical input (*36, 37*), whereas CGE-derived interneurons are recruited later via feedback from higher cortical areas and participate in modulatory or disinhibitory circuits (*38–40*). These differences are examined in greater detail below. Among non-neuronal types, oligodendrocytes were also more abundant in opossums than mice (Fig. 1K). Histological measurements confirmed elevated oligodendrocyte densities in adult opossums compared to those reported for ∼P40 mice, but comparable to P60 mice, consistent with age-dependent expansion of this population that continues into adulthood (fig. S1T; (*41*)).

### Intratelencephalic glutamatergic neurons have poor cross-species correspondence

While we successfully identified most glutamatergic, GABAergic, and non-neuronal subclasses in opossums, IT neurons posed a unique challenge. These neurons, which project within the cortex, are subdivided in mice and other eutherians into layer-specific populations based on gene expression (*35, 42*). In transcriptomic space, mouse IT neurons form a well-structured continuum (Fig. 2A–B, left) where unsupervised clustering and canonical markers (fig. S2A–D) resolve distinct subclasses in layers 2/3, 4, 5, and 6. Opossum IT neurons also formed a continuum (Fig. 2B, right) but lacked comparable organization and coherent expression of canonical layer markers (fig. S2H, L), precluding assignment to clear homologous subclasses. We therefore defined four opossum IT subclasses (IT_A–D) using unsupervised clustering (fig. S2E–M) and assessed their transcriptomic correspondence to the layer-specific IT types in the mouse.

**Figure 2.**
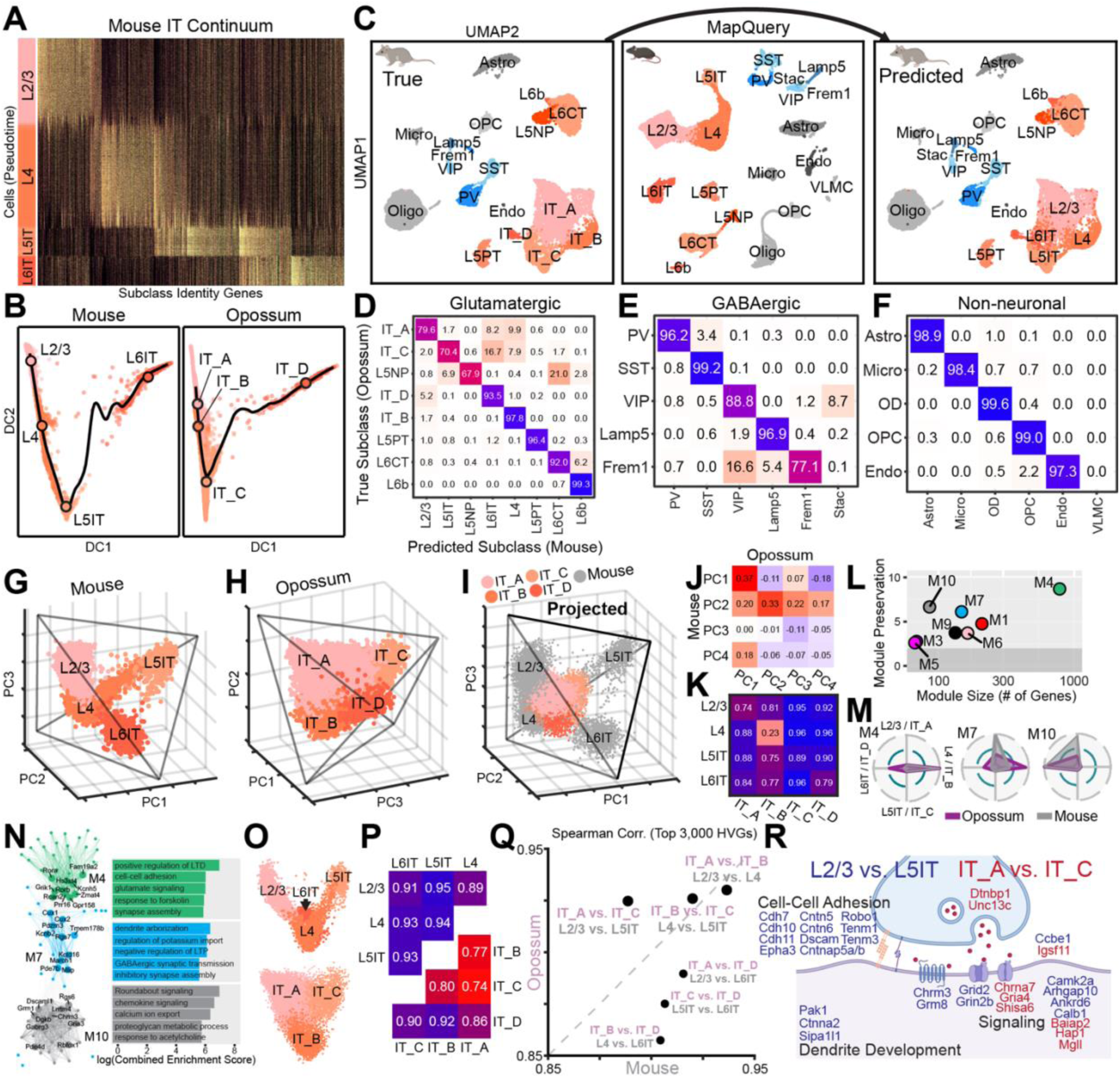
Intratelencephalic (IT) glutamatergic neurons have poor cross-species correspondence. (**A**) Mouse IT cells form a depth-ordered continuum (L2/3–L4–L5IT–L6IT) along a pseudotime axis derived from subclass identity genes, many of which are shared among spatially adjacent layer-specific populations. (**B**) Continuous variation between mouse (left) and opossum (right) IT subclasses in diffusion component (DC) space. Compared to mice, opossum IT cells display higher intermixing along the dominant trajectory (black line) and were labeled based on unsupervised clustering (see fig. S2). Black circles denote subclass means. (**C**) Opossum cells, shown by subclass in UMAP space (left) were assigned mouse subclass labels (right) using mouse cells (center) as a reference in Seurat’s MapQuery pipeline. (**D–F**) Confusion matrices (row-normalized) showing the relationship between true and predicted opossum subclass labels described in (C) for glutamatergic (D), GABAergic (E, and non-neuronal (F) subclasses, which were classified independently. (**G–H**) IT neurons in mice (G) and opossums (H) form tetrahedral gradients when projected onto their top three principal components (PCs) computed from full gene sets within each species. This organization is maintained in ortholog space (fig. S3A–B). Tetrahedrons were fit using a published Pareto task inference method. (**I**) Opossum IT neurons projected into mouse IT PC space. (**J**) Correlation matrix between the loadings (gene weights) of the top four within-species PCs from mouse and opossum IT neurons. (**K**) Average silhouette distances between cell embeddings of mouse and opossum IT subclasses in (I). (**L**) Weighted gene co-expression network analysis (WGCNA) modules identified in mouse IT neurons and projected onto opossum IT neurons, shown by size and cross-species correspondence (preservation, see Methods). (**M**) Subclass specificity of the three most highly preserved WGCNA modules (M4, M7, and M10) in mouse and opossums. (**N**) Top 20 hub genes associated with the modules in (M) shown as network diagrams and module-enriched gene ontology (GO) terms. (**O**) View of a single plane in species-specific mouse (top) and opossum (bottom) IT PC spaces containing gradients of corresponding subclasses. Opossum IT_D is fully obscured from this view. (**P**) Average silhouette distances between cell embeddings from mouse and opossum IT subclasses in species-specific (three-dimensional) PC spaces shown in (G–H). (**Q**) Spearman correlations between the top 3,000 highly variable genes (log-transformed expression levels, see also fig. S3E) in pairwise, within-species combinations of IT subclasses. Show as mouse vs. opossum against identity line to emphasize cross-species differences. (**R**) DEGs and enriched GO terms in mouse L2/3 vs. L5IT (blue) and opossum IT_A vs. IT_C (red) shown in their expected cellular compartment.

To do so, we restricted our analysis to genes with one-to-one orthologues (fig. S2N). This filtering had minimal impact on subclass separability within each species, which remained high (fig. S2O). We then applied Seurat’s MapQuery pipeline to project opossum cells into a mouse reference embedding based on gene expression and transfer labels using nearest neighbors (Fig. 2C). Some opossum subclasses mapped strongly onto mouse counterparts—for example, IT_B corresponded to thalamorecipient L4 and IT_D to L6IT. In contrast, IT_A, IT_C, and L5NP showed weak and inconsistent correspondence to any mouse IT subclass (Fig. 2D, fig. S2P). By comparison, other glutamatergic populations (e.g., L5PT, L6CT, and L6b), as well as most GABAergic and non-neuronal subclasses, showed strong cross-species alignment (Fig. 2E–F, fig. S2Q–R). Notably, the distinction between *Vip*- and *Frem1*-expressing interneurons—CGE-derived GABAergic subclasses that, like IT neurons, form a continuum—was substantially blurred across species (Fig. 2E). While this reflects the difficulty of defining ‘types’ within continuous cell populations, closer examination (described below) revealed fundamental differences between the structure of IT and CGE continua across species.

### Gene expression continua reveal conserved and divergent features of IT neurons

Gene expression continua, as observed for IT cells, can arise naturally during development but are also present in differentiated, multifunctional tissues that divide labor among competing functional demands. According to tissue multi-tasking theory (*29–31*), such division of labor is supported by cells that occupy Pareto fronts—low-dimensional structures in gene expression space that reflect optimal trade-offs between spatially restricted tasks. In the neocortex, division of labor is evident across layers: for example, neurons projecting to subcortical targets (e.g., L5PT, L6CT) have highly distinct transcriptomic profiles from IT neurons (*22, 35*). But within layer-specific IT populations, graded gene expression patterns give rise to low-dimensional structures that suggest partial functional overlap and division of labor among cell types (*43, 44*). Given the imperfect correspondence of IT subclasses between opossum and mouse, we examined the structure of their subclass-level IT continua in species-specific principal component (PC) spaces. Multi-tasking theory predicts that tissues with D-dimensional spatial zonation (here, D=4 cortical layers: 2/3, 4, 5, 6) will lie on a D-1-dimensional manifold in gene expression space (*29*). Indeed, both mouse and opossum IT neurons formed tetrahedral structures in the first three PCs (Fig. 2G–H, fig. S3A–B). Projecting opossum IT neurons into mouse PC space preserved this structure (Fig. 2I–J), but only L4 and IT_B showed substantial cross-species overlap (Fig. 2K), consistent with their unique cross-species correspondence discussed above.

Moreover, the mouse IT-specific weighted gene co-expression network analysis (WGCNA) module with the greatest cross-species preservation was large (>500 genes) and L4-specific (Fig. 2L–M). Other conserved modules (M7 and M10) were shared across L4–IT_B and L6IT–IT_D but were weakly represented in opossum IT_A despite strong L2/3 representations in mice (Fig. 2M), highlighting the divergence of upper-layer IT neurons across species. Gene ontology analysis of the conserved M4 and M7 revealed an enrichment for genes regulating synaptic plasticity, cell-cell adhesion, synapse assembly, and dendritic arborization, suggesting conserved functional and anatomical properties in the thalamorecipient layer (Fig. 2N).

Principal component analysis further revealed that while mouse IT cells varied continuously between L4 and other IT subclasses, L2/3, L5IT, and L6IT were more discrete and occupied distinct vertices of the tetrahedron (Fig. 2G, O, fig. S3A). By contrast, in opossum, the tetrahedral IT structure comprised a triangular plane containing a continuous gradient of IT_A, IT_B, and IT_C cells (Fig. 2O–P, fig. S3B), with IT_D cells partially overlapping with the L4-like IT_B (fig. S3B, bottom left). Importantly, this gradient phenomenon is unique to the IT continua (fig. S3C–E) and does not arise from low-quality or noisy gene expression data in opossums (fig. S3F–H, see Methods). In line with the unique variation between IT_A and IT_C in opossums (Fig. 2O), the correlation of their top highly variable genes was far higher than mouse L2/3 and L5IT (Fig. 2Q, see also fig. S3E). The top within-species DEGs between these subclasses were enriched for genes involved in neuronal signaling, including ionotropic receptor subunits, and synaptic release (Fig. 2R). However, mouse-specific DEGs (between L2/3 and L5IT) were also enriched for genes related to adhesion and dendritic development, suggesting additional specialization in connectivity and morphology. Together, these results suggest that opossum IT neurons exhibit more generalized transcriptomic identities than mouse IT neurons, particularly along the IT_A–IT_C (L2/3–L5IT) axis.

### A spatial gradient along the opossum IT_A–C transcriptomic axis

When applied to tissues, Pareto optimality predicts that cells that are transcriptomically specialized—like those at the vertices of a tetrahedron in PC space—should also be spatially segregated (*29*). Conversely, generalist continua are theorized to emerge under conditions where spatial gradients are present in tissue. Given the continuous nature of opossum IT neurons, we asked whether their laminar organization differed from mice. To do this, we performed whole-transcriptome spatial profiling of mouse and opossum V1 using Stereo-seq (*45*).

Following multi-embedding of coronal sections and the Stereo-seq pipeline (see Methods), we identified layers 1–6 within V1 using anatomical landmarks and canonical marker genes (Fig. 3A–B), extracted cortical columns from multiple sections (fig. S4K–L), and assigned cell subclass identities via co-clustering with snRNA-seq atlases (Fig. 3A, C, fig. S4E–J; subclasses with less than 25 total cells were excluded from downstream analyses). Subpial density profiles revealed largely consistent spatial organization in glutamatergic subclasses from both species (Fig. 3D). Notably, opossum IT_A displayed a somewhat broader distribution than mouse L2/3 (Fig. 3D, left), but other IT and non-IT subclasses were well-organized (Fig. 3D). GABAergic and non-neuronal subclasses largely followed expected depth distributions (fig. S4N–O).

**Figure 3.**
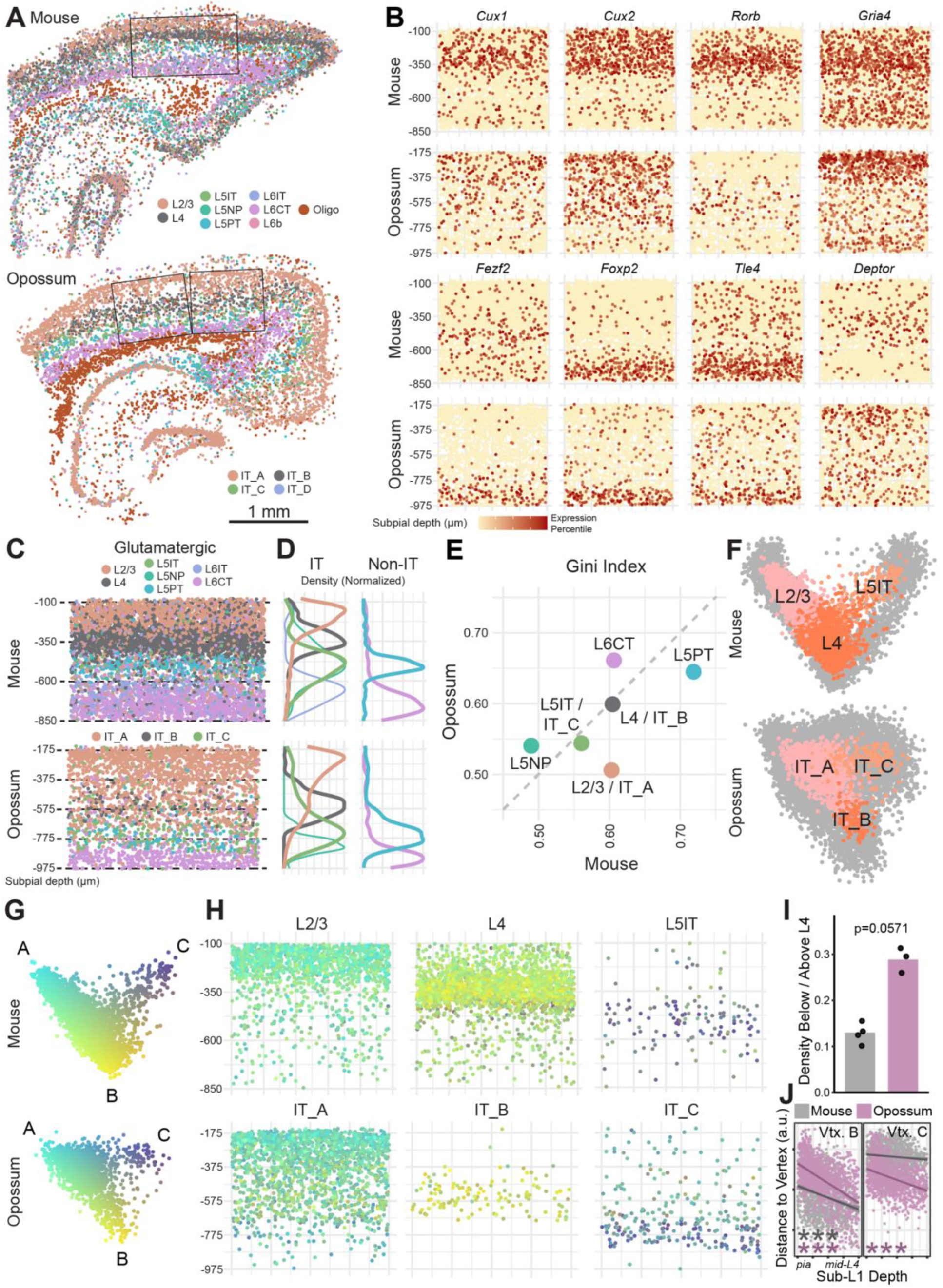
Stereo-seq reveals different spatio-transcriptomic axes in opossums and mice. (**A**) Representative Stereo-seq sections from mouse (top) and opossum (bottom), displaying all glutamatergic subclasses and oligodendrocytes. Subclass labels were predicted from species-specific snRNA-seq atlases using integration and co-clustering (see Methods). V1 was identified using cytoarchitectural features and the prominent presence of L4–IT_B. Boxes indicate regions of interest (ROIs) from these sections used for downstream analyses. (**B**) Spatially variable genes include canonical upper- and lower-layer glutamatergic subclass markers. (**C**) Distribution of glutamatergic subclasses from pooled V1 regions. (**D**) Subpial densities of IT (left) and non-IT (right) glutamatergic subclasses, collapsed along the x-axis in (C). (**E**) Gini index of subpial position for glutamatergic subclasses in mice and opossums. (**F**) IT cells identified in Stereo-seq (colored) shown against snRNA-seq IT cells (gray) in pooled PC space (similar to Fig. 2O). (**G**) Stereo-seq IT cells in PC space as in (F), pseudocolored by distance from the vertices of a fitted triangle. (**H**) Pseudocolored cells from (G) shown in physical space as in (C), grouped by subclass labels shown in (F). (**I**) Ratio of densities of L2/3 (mouse) and IT_A (opossum) cells below and above the species-specific median subpial depth of L4–IT_B (Wilcoxon rank-sum test; p=0.0571; table S1). Opossum data points include three ROIs from two sections, and mouse data points include four ROIs from three sections. (**J**) Relationships between the depth of L2/3 (mouse) and IT_A (opossum) cells and their distance from B (left) and C (right) vertices in PC space (**G**). The depth axis extends from the bottom of L1 to the middle of L4–IT_B in each species. Lines are ordinary least-squares fits. Significance markers reflect partial Spearman correlations along each transcriptomic axis, accounting for covariation of distance from B and C vertices (***p<0.001; table S1).

To quantify spatial organization, we computed Gini indices from subpial depth distributions of each glutamatergic subclass and plotted cross-species comparisons (Fig. 3E). This measures the degree to which cells are restricted to a single depth, with higher values indicating stronger laminar confinement. Subclasses with input and output roles—L4–IT_B, L5PT and L6CT—showed the highest spatial organization in both species, consistent with the notion that these subclasses are the most tightly conserved.

To further explore the relationship between spatial and transcriptomic variation in IT neurons, we integrated Stereo-seq and snRNA-seq data from L2/3–IT_A, L4–IT_B, and L5IT–IT_C cells and projected them onto shared principal components (Fig. 3F). In both species, Stereo-seq cells formed triangular gradients in PC space, reminiscent of our snRNA-seq findings (Fig. 2O). We then pseudocolored Stereo-seq IT cells in the gradient by their distance from each of three triangle vertices—A (L2/3-like), B (L4-like), and C (L5IT-like)—defined in PC2-PC3 space (Fig. 3G). Spatial mapping of these pseudocolored cells emphasized the broad distribution of IT_A cells, which were denser below L4 than L2/3 cells (Fig. 3H, I). We also observed that IT_A cells closest to transcriptomic vertex A (bright cyan) were concentrated in putative layer 2/3 (Fig. 3H). In mice, L2/3 cells show a continuous gradient toward L4 in both transcriptomic and physical space, such that deeper L2/3 cells have gene expression profiles more similar to L4 cells (Fig. 3J; (*35, 43, 44*). This relationship also holds true for opossum IT_A, which varies significantly with depth relative to vertex B. Critically, opossum IT_A varies similarly with respect to vertex C (L5IT-like), a pattern absent in mouse L2/3 (Fig. 3J; see Methods). This unique spatio-transcriptomic IT_A–C axis provides strong evidence that opossum IT neurons—particularly IT_A—exhibit generalist identities.

### Divergent density and perineuronal net association of PV neurons across species

To validate the cross-species differences in GABAergic subclass proportions observed in our transcriptomic data (Fig. 1J), we performed immunofluorescent staining in mouse and opossum V1. Consistent with our snRNA-seq findings, the density of PV-positive neurons was ∼20% higher in opossums than in mice (Fig. 4A, B), with the most pronounced difference in layers 4 and 5—the layers with the highest PV density in both species. Nonetheless, PV neuron densities were broadly scaled across layers in opossums (Fig. 4C). These differences were not explained by global differences in neuronal packing density (Fig. 4D, E), though NeuN staining revealed that layer 6 is significantly denser in opossums than mice.

**Figure 4.**
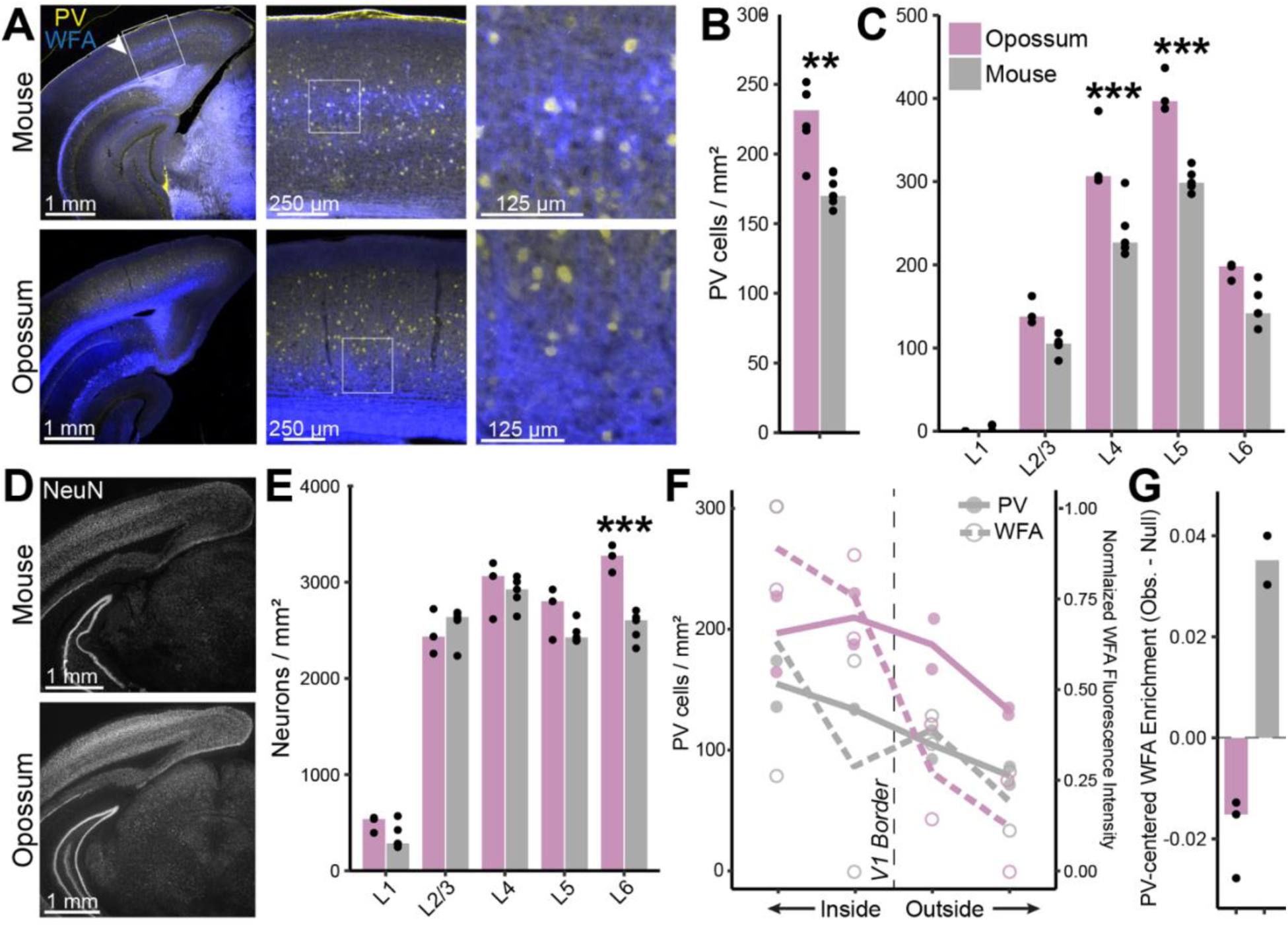
Immunolabeling confirms cross-species differences in PV density and peri-neuronal net localization. (**A**) Parvalbumin (PV; blue) and Wisteria floribunda agglutinin (WFA; yellow) labeling in representative mouse and opossum coronal sections. Arrows mark the lateral edge of V1, and boxes indicate the magnified area shown on the right. (**B**) PV-positive cell densities in V1 are significantly different (Wilcoxon rank-sum test; *p < 0.05, **p < 0.01, ***p < 0.001; table S1; data points are single fields of view from separate sections). Opossum and mouse data points include five sections from each species. (**C**) layer-specific PV cell densities (identified cytoarchitecturally with NeuN counterstaining) are significantly different in layers 4 and 5 (ANOVA followed by two-sided Tukey’s HSD test and Bonferroni correction; table S1). (**D**) NeuN labeling. (**E**) Layer-specific neuron densities in V1 reveal significantly higher packing density in opossum layer 6 (ANOVA followed by two-sided Tukey’s HSD test and Bonferroni correction; table S1). Opossum and mouse data points include three and five sections, respectively. (**F**) PV density and WFA fluorescence intensity in bins located inside and outside of V1 (see Methods). (**G**) Median WFA fluorescence in an annulus surrounding PV-positive neurons compared to a depth-matched null distribution sampled from random nearby points (see Methods). Bars show the median difference between observed and null values. Positive values indicate preferential WFA enrichment around PV neurons.

Given the established role of PV neurons in regulating critical period plasticity (*32, 46*), we examined the distribution of perineuronal nets (PNNs) using Wisteria floribunda agglutinin (WFA) staining (Fig. 4A). In mouse V1, WFA staining was densest in layer 4, consistent with previous reports showing that PNNs are enriched in thalamorecipient layers and frequently surround PV neurons, where their density correlates with thalamic input strength (*47*). By contrast, WFA staining was largely restricted to deeper layers 5b and 6 in opossums (Fig. 4A), in agreement with a previous report indicating that PNNs preferentially surround layer 5 pyramidal neurons in this species (*48*). Outside of V1, both PV neuron density and WFA fluorescence declined in both mice and opossums (Fig. 4A, F).

To test whether PNNs are preferentially localized around PV neurons, we quantified WFA fluorescence in an annulus surrounding PV-positive cells and compared it to a depth-matched null distribution sampled from nearby locations (Fig. 4G; see Methods). In mouse V1, WFA fluorescence was enriched around PV neurons relative to the null distribution, indicating a specific PV–PNN association. By contrast, opossum PV neurons showed no such enrichment; observed WFA signal did not exceed the null distribution, consistent with the absence of PV-centered PNN localization.

## Discussion

Metatherian and eutherian mammals diverged roughly 160 to 180 million years ago during the Middle to Late Jurassic, providing a deep phylogenetic contrast for examining variation in mammalian cortical organization (*13–15*). Our data reveal substantial differences in the cellular and spatial organization of the neocortex between the gray short-tailed opossum (*Monodelphis domestica*) and the laboratory mouse (*Mus musculus*). Importantly, these differences do not reflect ancestral cortical states but instead highlight specific cellular populations whose organization differs across these deeply diverged mammalian lineages.

The most prominent differences we observed involve intratelencephalic (IT) glutamatergic neurons, particularly L2/3 and L5IT subclasses. In mouse cortex, these populations exhibit greater transcriptomic differentiation and sharper separation in gene expression space, consistent with increased specialization of intracortical communication pathways. This pattern aligns with theoretical frameworks in which cell types diversify along Pareto fronts, with specialization emerging through trade-offs between competing functional demands (*29–31*). In contrast, opossum IT neurons display broader and more overlapping transcriptomic profiles, suggesting more generalized functional identities within intracortical circuits. While the specific computational tasks driving these differences remain unknown, they likely relate to processes such as sensory integration, inter-areal communication, and experience-dependent modulation.

The spatial organization of IT neurons supports this interpretation. In both species, IT populations exhibit structured laminar distributions, indicating that spatial organization is largely preserved across lineages. However, these distributions differ in precision. In mouse cortex, the layer 2/3 IT population is more sharply confined to its discrete laminar zone above layer 4, while opossum IT_A occupies a broader laminar range. By contrast, thalamorecipient L4 neurons, extratelencephalic L5PT neurons, and corticothalamic L6 neurons remain remarkably conserved in both transcriptomic identity and spatial organization across species, suggesting that evolution may have disproportionately affected intracortical rather than input–output pathways. This pattern contrasts with the retina, where evolutionary divergence is greatest among output ganglion cells (*28*). Crucially, we also demonstrated the existence of a unique spatio-transcriptomic axis between IT_A and IT_C in opossums, supporting the notion that opossum IT neurons along this axis have generalist identities (*29*).

We also uncovered intriguing differences within populations of inhibitory neurons. Compared to opossums, mice show a shift in inhibitory composition away from PV interneurons toward CGE-derived subclasses, including VIP cells. This pattern may reflect enhanced flexibility, disinhibition, and neuromodulation, reducing emphasis on rigid temporal precision and promoting adaptable cortical computations. PV interneurons receive robust feedforward excitation and provide perisomatic intracortical inhibition that sharpens spike timing and enhances gamma oscillations (*36, 49, 50*). The relative expansion of CGE-derived interneurons in mice may therefore reflect greater reliance on flexible, state-dependent cortical modulation (*51–53*), whereas stronger perisomatic inhibition in opossums may favor more feedforward-driven processing. Although interneuron proportions vary across mammalian species (*22*), molecular differences within excitatory IT populations appear more pronounced than those among inhibitory classes, even when including primates such as marmosets and humans (*22, 54–56*).

Additional differences in plasticity-associated features reinforce this distinction. Perineuronal nets (PNNs) preferentially surround PV interneurons in mice but not opossums, where they appear to be associated with L5 pyramidal neurons (*48*). In eutherian species, PNNs stabilize inhibitory circuits and contribute to critical period closure and memory consolidation (*33, 34*). Their association with pyramidal neurons in opossums suggests that plasticity constraints are imposed directly on cortical output pathways, while in mice, PNN localization places these constraints upstream, on inhibitory circuits that gate thalamocortical input.

Together, these findings demonstrate that while the basic laminar framework and major neuronal subclasses of the cortical column are conserved, substantial variation exists in the specialization, spatial organization, and plasticity-related organization of intracortical circuits across mammalian lineages. IT neurons emerge as a principal axis of divergence, providing a comparative foundation for future studies aimed at understanding how cortical specialization and adaptability vary across mammals.

## Methods

### Animals

Mouse procedures were performed under IACUC protocol ARC-2009-031 at the University of California, Los Angeles. One adult (∼P40) male wild-type C57BL/6J mouse was used for spatial transcriptomics. Two additional adult (∼P40) males were used for immunofluorescence. Single-nucleus RNA sequencing data from adult mice (P38) was obtained from our previous study (*35*).

Opossum (*Monodelphis domestica*) procedures were also performed under ARC-2009-031. Two adult (1.5–2 years) male opossums were used for single-nucleus RNA sequencing, one younger adult (∼6 months) male opossum was used for spatial transcriptomics, and two additional adult (1.5–2 years) males were used for immunofluorescence.

### V1 dissection

Opossums were deeply anesthetized with isoflurane followed by decapitation, and brains were removed into ice-cold Hibernate A (BrainBits Cat. # HACA). Anatomical landmarks (*20*) were used to identify area 17 (V1). A 2 mm coronal slice was obtained by bisecting the superior colliculus and making a second cut 2 mm anterior to the first. Cuts were also made lateral to V1 (∼3 mm from the midline) in both hemispheres. V1 was dissected from the underlying white matter, flash-frozen in liquid nitrogen, and stored in liquid nitrogen until single-nucleus RNA sequencing (∼1 year).

### Nuclei isolation and droplet-based single-nucleus RNA sequencing

Nuclei were isolated from each biological replicate using the 10x Genomics Chromium Nuclei Isolation Kit. We performed fluorescence-aided nuclear sorting (FANS) for Draq5-positive nuclei at the UCLA Flow Cytometry Core. After FANS, nuclei suspensions were counted on a hemocytometer using an ethidium bromide counterstain and diluted to 700-1200 cells per µL. Nuclei suspensions from each biological replicate were loaded into two wells of a 10x Chromium 3’ v3 chip, each targeting 10,000 cells. We refer to these as library replicates. Libraries were pooled into a single lane of an Illumina NovaSeq X Plus 10B and sequenced to ∼30,000 reads (2×50 bp) per cell. Library preparation and sequencing were performed by the UCLA Technology Center for Genomics and Bioinformatics (TCGB).

### Spatial transcriptomics (Stereo-seq v1.3)

Animals were deeply anesthetized with isoflurane followed by decapitation, and brains were immediately removed and submerged in ice-cold 1x PBS containing RNase inhibitor (1 unit per µL; Thermo Fisher Cat. # AM2696). Extracted brains were placed in an ice-cold metal mold and, in the opossum, 2 mm coronal slices were obtained by bisecting the superior colliculus with one blade and inserting additional blades 2 mm anterior and posterior to the first (*20*). In the mouse, the first blade was inserted at the cortical margins of the midline (below the lambdoid suture) with additional blades 1 mm anterior and posterior. Slices were placed back into ice-cold PBS. To embed two coronal slices (containing four total V1 hemispheres) in the opossum, cuts were made lateral to V1 (∼3 mm from the midline) in both hemispheres, as well as horizontal (∼4 mm from the highest point of cortical surface). Because of the difference in brain size, these cuts were not necessary in the mouse. Coronal slices were arranged in an ice-cold base mold containing a drop of OCT (Tissue-Tek), then covered in OCT and flash-frozen in liquid nitrogen. Blocks were stored at −80°C and sectioned at −20°C. Stereo-seq was performed according to Complete Genomics user manuals. Sectioning and imaging were performed by the UCLA Translational Pathology Core Laboratory (TPCL). RNA integrity (RIN) analysis, permeabilization, library preparation, and sequencing was performed by the UCLA TCGB.

RNA integrity numbers (RINs) obtained from block shavings were 7.1 for opossum tissue and >9 for mouse tissue, both above the recommended threshold of 4.0 for Stereo-seq v1.3. Permeabilization time was optimized according to vendor guidance and set to 12 minutes for both mouse and opossum samples based on maximal fluorescence signal with minimal diffusion. For transcriptomics, a single 10 µm section was cut and mounted onto the 10 mm x 10 mm chip on the Stereo-seq Chip T slide, fixed in pre-cooled methanol (30 minutes), and processed with 12-minute tissue permeabilization before in situ reverse transcription. cDNAs were then released, denatured, amplified, and purified, and libraries were sequenced on a full MGI DNBSEQ-T7 flow cell with paired-end 75 bp reads. Stereo-seq data processing is discussed in detail below.

### Immunofluorescence

Animals were deeply anesthetized with isoflurane followed by decapitation. Brains were removed into ice-cold 1x PBS (5 min), transferred to ice-cold 4% paraformaldehyde (PFA; 5 min), post-fixed overnight at 4°C, and cryoprotected in 20% sucrose before embedding in OCT. Blocks were stored at −80°C until cryosectioning at −20°C to obtain 40 µm free-floating coronal sections spanning the rostrocaudal extent of V1. Sections were permeabilized and blocked (0.2% Triton X-100 and 10% normal goat serum in 1x PBS) for 1 hour at room temperature (RT), incubated in primary antibodies overnight at 4°C, washed (3×10 min in 1x PBS), incubated in secondary antibodies (2 h, RT), and washed again (3×5 min) before imaging. Primary antibodies and concentrations used were as follows: mouse anti-PV (1:1000; Millipore Sigma Cat. # P3088), wisteria floribunda agglutinin (WFA; 1:500; Vector Laboratories Cat. # FL-1351-2), mouse anti-CC1 (1:200; Millipore Sigma Cat. # OP80), rat anti-myelin basic protein (MBP; 1:200; Bio-Rad Cat. # MCA409S), and rabbit anti-NeuN (1:1000; Abcam Cat. # ab177487). Secondary antibodies and concentrations used were as follows: goat anti-mouse Alexa Fluor™ 594 (1:1000; Thermo Fisher Cat. # A-11005), goat anti-rabbit Alexa Fluor™ 488 (1:1000; Thermo Fisher Cat. # A-11008), and goat anti-rat Alexa Fluor™ 488 (1:1000; Thermo Fisher Cat. # A-11006).

### Microscopy

Sections were mounted on glass slides and coverslipped with antifade mounting media (Vector Laboratories Cat. # H-1000-10). Anatomical landmarks were used to identify sections containing V1. Images were acquired on an Olympus BX51WI Upright Fluorescence Microscope equipped with a Prior Lumen 200 Fluorescence Illuminator and a Basler Ace Classic camera (acA2040-90um). Multiple sections from each animal were imaged, with medial and lateral fields collected from each hemisphere at 4x where possible.

### Analysis of single-nucleus transcriptomics data

Mouse snRNA-seq data (*35*) were obtained from Gene Expression Omnibus (GSE190940; P38 normally reared samples GSM5754081–3). Raw counts were matched with metadata (including ‘Class’, ‘Subclass’, and ‘Type’) from processed H5AD objects (https://github.com/-shekharlab/mouseVC). These data comprised two biological replicates, one split into two library replicates. Seurat v4.3.0 (*57*) objects were constructed and processed identically to opossum snRNA-seq data (discussed in detail below), without gene ortholog mapping or class/subclass assignment.

Raw reads (FASTQ) from opossum snRNA-seq libraries were aligned to the *Monodelphis domestica* reference genome ASM229v1 (*58*) with Cell Ranger v6.1.1 on the UCLA Hoffman2 cluster with --include-introns set to true. The reference package was generated from Ensembl Release 110 FASTA/GTF files using Cell Ranger’s mkref function. Despite similar sequencing depths (fig. S1A, top), opossum alignments showed substantially lower ‘reads mapping confidently to transcriptome’ (fig. S1A, bottom), and considerably more intergenic reads (mapping confidently to regions outside of exons, introns, or UTRs). Examination of the reference and alignments revealed many genes that contained only coding sequences (often predicted) and lacked introns and 5’ or 3’ untranslated regions (UTRs), as well as prominent intergenic read pileups directly downstream (3’) of the annotation. We quantified intergenic reads and their proximity to the nearest same-strand gene using Samtools and BEDtools (fig. S1B), stratifying by gene biotype and 3’ UTR annotation. This revealed a clear genome-wide pileup within 5 kbp downstream of the nearest gene for protein-coding genes with or without 3’ UTRs, along with additional peaks downstream from long noncoding RNAs (lncRNAs). In mice—where most genes have an annotated 3’ UTR (fig. S1C, inset)—a similar downstream signal was present but constituted a lower fraction of total reads. Notably, the downstream intergenic read distribution for opossum genes with a 3’ UTR was bimodal. Consistent with this, annotated 3’ UTRs were broadly shorter in the opossum genome (fig. S1C) and showed spikes at 3 bp (stop codons) and 1000 bp, suggestive of annotation artifacts that could shift apparent pileups further downstream. In light of these artifacts and the presence of downstream pileups for other biotypes, we extended 3’ UTRs up to 3 or 5 kbp downstream from the 3’ end of annotation for every gene farther than 10 kbp upstream of the nearest same-strand gene (fig. S1D) and re-aligned reads to this custom genome. This resulted in a ∼5-10% increase in genes and transcripts (unique molecular identifiers, UMIs) per cell (fig. S1E–F). Although these metrics remained considerably lower than mouse data (fig. S1E–F), 3’ extension increased cross-species class-specific pseudobulk expression correlations while minimally affecting the manifold (fig. S1G), particularly between the 3 and 5 kbp extensions. We therefore performed all downstream analyses, including spatial transcriptomics, on data aligned to the genome containing custom 5-kbp 3’ UTRs.

All analyses were performed in R (v4.3.0) and using Seurat (v4.3.0) unless otherwise specified. Gene expression matrices from Cell Ranger were imported into Seurat. Opossum data comprised four samples (two biological replicates, each split into two library replicates). As in (*35*), we filtered cells (700 < genes < 6500 and UMIs < 40,000) and removed genes expressed in 8 or fewer cells. Because mitochondrial genes are not annotated in the opossum genome, we did not filter by mitochondrial UMI fraction. This yielded 32,764 cells and 26,440 genes. Opossum gene names were mapped to mouse gene names when a one-to-one ortholog existed (Ensembl BioMart). Unless otherwise specified, within-species analyses used full gene sets, including species-specific genes.

Cells were normalized using Seurat’s SCTransform (v2) workflow (*59*). We computed 30 principal components (PCs), built a nearest-neighbor graph, clustered using the Leiden algorithm (resolution = 1) and visualized with uniform manifold approximation (UMAP). In opossums, this yielded 27 initial clusters (fig. S1H), with representation from all samples (fig. S1I–J) and less than 15% predicted doublets per cluster (fig. S1K) identified using Scrublet (*60*). Clusters were divided into glutamatergic, GABAergic, and non-neuronal classes using canonical marker genes (fig. S1L–M), then each class was re-normalized and re-clustered (fig. S1N–P). Based on cross-species correspondence (label transfer, see below), predicted doublet composition, or size (fig. S1Q–S), seven GABAergic clusters (10, 11, 12, 14, 15, 17, and 18) and five non-neuronal clusters (7, 10, 11, 14, 16) were excluded from downstream analyses.

Within each class, we re-normalized and re-clustered at Leiden resolutions 0.5, 1, and 2 and assigned subclasses based on canonical markers (Fig. 1E). Opossums lacked homologs of the mouse GABAergic subclass Stac and non-neuronal vascular and leptomeningeal cells (VLMC). As discussed further below, because glutamatergic intratelencephalic (IT) neurons displayed ambiguous marker expression, we defined 4 opossum IT subclasses (IT_A–D; matching the number of mouse IT subclasses) with unsupervised clustering (fig. S2E–M). Subclass assignments were evaluated using 1) within-species label transfer (MapQuery) and 2) differential gene expression analysis (see below). These analyses yielded measures of subclass consistency and separability in opossums that reflected our observations in mice. Cell types were labeled in a similar fashion but are not discussed due to limited interpretability in opossums.

For cross-species or cross-modal integration (Fig. 1F, 3F, fig. S4E–J), Seurat objects were subset to shared genes (one-to-one orthologs for cross-species), then normalized and clustered in that shared space. We applied SCTransform integration (SelectIntegrationFeatures, PrepSCTIntegration, FindIntegrationAnchors, and IntegrateData) with normalization.method = “SCT”, and features.to.integrate set to shared genes. Integrated objects were visualized using PCA, nearest-neighbor graph construction/clustering, and UMAP. Fractional subclass overlap in the integrated space (Fig. 1F) was computed by summing, across integrated clusters, the minimum fraction of each subclass pair within each cluster.

Within- and cross-species label transfer was performed using Seurat’s MapQuery (fig. S1Q–S, Fig. 2C–F, fig. S2C, F–G, J–K, O). For within-species transfer (fig. S2C, F, G, O), cells were randomly divided into two objects (full gene sets, except for fig. S2O), normalized and clustered, then used as a reference for label transfer to the other (query) object; predicted labels and probabilities were pooled across splits. For cross-species transfer (Fig. 2C, fig. S2G, K), genes were subset to one-to-one orthologs prior to normalization and clustering, with mouse as the reference. To balance subclasses, we performed 100 iterations in which each subclass contributed at most 100 randomly sampled cells; cells were depleted across iterations until exhausted, then the pool was reset. Per iteration, FindTransferAnchors used reference.reduction = “pca”, dims = 1:30, and MapQuery used reduction.model = “umap”. Confusion matrices comparing predicted (rows) and true (columns) labels were row-normalized.

IT neurons were examined species-specific PC spaces (Fig. 2G–H). PCs were computed from full gene sets, but their overall structure was preserved in ortholog space (Fig. 2I, fig. S3A–B). To account for subclass imbalance, each IT subclass was downsampled to the least common (L6IT/IT_D) for PC computation, and all cells were projected onto these PCs. Pareto Task Inference (*31*) (ParTI; MATLAB r2023a; https://github.com/AlonLabWIS/ParTI) was used to fit polytopes to IT cells using PCs 1–30; optimal vertex number (2–10) was evaluated with the principal convex hull algorithm (PCHA) and selected with the elbow method, yielding tetrahedra (n = 4) in each species. Tetrahedron vertices were projected into PC1–3 space (Fig. 2G–I, fig. S3A–B). We computed pairwise correlations between gene loadings of the top 4 within-species PCs in mice and opossums (Fig. 2J) by subsetting one-to-one orthologs from the loadings vectors. Using a previously established shuffling procedure (*43*), we tested whether opossum and mouse IT cells form a genuine continuum and are not discrete subclasses that appear continuous due to noisy gene expression (fig. S3F–G). We also tested whether cross-species differences in IT continua were attributable to differences in sequencing depth and transcriptome coverage (fig. S1A, E, F, S3H) by downsampling the mouse UMI count distribution to match the opossum. For each mouse cell, we randomly sampled a target UMI count from the empirical distribution of total UMI counts observed in opossum. Mouse gene expression profiles were then downsampled using multinomial resampling: for each cell, gene-level counts were proportionally reduced to the sampled target UMI total while preserving the original relative expression profile within the cell. This procedure was implemented using the raw count matrix and performed independently for each cell.

We applied high-dimensional weighted gene co-expression network analysis (hdWGCNA) (*61*) (https://smorabit.github.io/hdWGCNA) to opossum and mouse IT cells (Fig. 2L–N). The standard pipeline identified 10 modules in mouse IT cells; after subsetting to one-to-one orthologs, modules were projected onto opossum IT cells with ProjectModules and evaluated with ModulePreservation. Modules with fewer than 50% one-to-one orthologs (M2) or with extremely poor preservation (M8) were excluded. We visualized subclass-specific preservation of modules (Fig. 2M) using ModuleRadarPlot and hub genes from highly preserved modules (Fig. 2N) using HubGeneNetworkPlot. Gene ontology (GO) was performed and visualized (Fig. 2N) with RunEnrichr and EnrichrBarPlot, respectively.

In Fig. 2Q, we computed within-species Spearman correlations between subclass-average expression profiles using the top 3,000 highly variable genes (HVGs; FindVariableFeatures) from log-normalized UMI counts. In fig. S3E, we z-scored log-normalized UMI counts within IT of CGE populations before computing these correlations. In these visualizations, each point represents a homologous pair of mouse (e.g., L2/3 vs. L5IT) and opossum (e.g., IT_A vs. IT_C) correlations.

Differential expression (Fig. 2R) was performed with FindMarkers using log2FC > 0.75, with pct.1 > 0.25 and p < 0.05 for IT_A vs. IT_C and L2/3 vs. L5IT comparisons. GO analysis was performed using GOfuncR (https://github.com/sgrote/GOfuncR) restricted to orthologous DEGs and the *Mus musculus* database available on Bioconductor (see go_enrich). We considered enriched ‘biological process’ terms with family-wise error rates less than 0.05. In Fig. 2R, we summarize top GO biological processes and display several associated genes for opossum- and mouse-specific comparisons.

### Analysis of spatial transcriptomics data

Spatial GEMs were generated from Stereo-seq FASTQ files using the Stereo-seq Analysis Workflow (SAW v7.1.2) with manual tissue alignment. Opossum reads were aligned to the same custom genome used for snRNA-seq, and mouse Stereo-seq reads were aligned to GRCm39 (GCA_000001635.9). We used the cellbin.adjusted SAW output format. Cells with fewer than 50 detected genes or fewer than 100 UMIs were removed (fig. S4A), as were genes expressed in 8 or fewer cells. One micron corresponds to two spatial units in Stereo-seq output (Complete Genomics). Absolute or relative cell densities were not quantified with Stereo-seq due to transcript dropout potential, which may be more pronounced in opossums (fig. S4A, M).

V1 regions of interest (ROIs) were manually selected using anatomical landmarks in both species (Fig. 3A, fig. S4K–L). ROIs extended from the top of layer 2/3 to the bottom of layer 6. Due to cortical curvature, multiple adjacent ROIs (cortical columns) were selected in some sections. Within each species, ROIs were combined into a ‘pseudocolumn’ by scaling each ROI to the width and height of the largest ROI. Column heights required minimal scaling (fig. S4K–L), and gene expression patterns and subclass distributions were consistent along ROI width (Fig. 3B, fig. S4K–L).

Clustering Stereo-seq pseudocolumns in transcriptomic space yielded little structure (fig. S4B–C), prompting integration with snRNA-seq atlases within each species (fig. S4E–J). Due to a lack of co-clustering using the standard Leiden algorithm, labels were assigned to Stereo-seq cells using k-nearest neighbor classification in integrated PCA space (top 30 PCs), assigning labels only when a sufficient percentage of their cross-modal (snRNA-seq) neighbors belonged to a single class or subclass.

Class assignment was performed over three rounds using agreement thresholds of 100%, 100%, and 75% among 50 nearest sRNA-seq neighbors (fig. S4E–G); cells labeled after rounds 1 or 2 were removed before the subsequent round, and cells failing all rounds were excluded (<10% in both species). Subclass assignment followed an analogous three-round procedure with thresholds of 100%, 75%, and 50% (fig. S4H–J). In the first round of each procedure (fig. S4E, H), snRNA-seq cells were randomly downsampled to match Stereo-seq cell numbers. Resulting subclasses closely matched expected spatial distributions (Fig. 3A, C–D, fig. S4K–L, N–P). Subpial density profiles were computed using geom_density.

To assess whether cortical depth independently correlated with transcriptomic distance to IT subtype vertices (Fig. 3J), we computed partial Spearman correlations controlling for distance to the alternate vertex using the ppcor package. Statistical significance was assessed by permutation testing (10,000 iterations). In mice, the partial correlation between depth and distance to the L5IT vertex became negative when controlling for distance to L4, consistent with an ordered L2/3–L4–L5IT trajectory.

In opossum snRNA-seq data, IT_A comprised three Leiden clusters (resolution 0.3) combined during subclass assignment (fig. S2I–M). Cluster labels were transferred to Stereo-seq pseudocolumn cells using nearest-neighbor assignment. Two major IT_A subclusters (1 and 2) were broadly distributed, while a smaller subcluster (5) that peaked solely in the deeper layers appeared unlikely to drive spatial effects (fig. S4P, left, dashed line is normalized to the peak of cluster 1). Despite IT_A subcluster 1 showing enrichment for species-specific lncRNAs (fig. S4Q), its distribution was closely aligned with subcluster 2. All IT_A subclusters have the highest gene expression correlations with mouse L2/3 (fig. S4R), with graded spatial and transcriptomic relationships to L5IT examined in Fig. 3G–J. Thus, IT_A subclusters could be conceptualized as cell types with differing spatial arrangements, as in mouse L2/3 (*35, 43, 44*).

To visualize transcriptomic gradients in Stereo-seq cells, IT subclasses (L2/3–IT_A, L4– IT_B, and L5IT–IT_C) were projected into PC space (Fig. 3F). Triangles were fit using a Python (v3.9.19) implementation of PCHA (https://github.com/ulfaslak/py_pcha; noc = 3, delta = 0.1), and cells were colored using a barycentric interpolation scheme based on their vertex proximity (Fig. 3G–H).

### Analysis of immunofluorescence data

Automated segmentation of PV- and NeuN-positive cells was performed using Cellpose (*62*) with the cyto2 model (cell diameter = 8, flow threshold = 0.5, cell probability threshold = −2.5), followed by manual curation. All analyses used mask centroids and were summarized at the section (hemisphere) level. Layer-specific PV cells densities (Fig. 4C) were obtained by manual selection of the neocortical layers using the NeuN channel.

For WFA analyses, images were background-subtracted using morphological opening (structuring element radius ∼50 µm) and intensity-normalized to the 99.5th percentile of cortical WFA signal. Cortical boundaries, the pial surface, and the lateral extent of V1 were manually annotated for each section.

To quantify areal variation in PV density and WFA fluorescence (Fig. 4F), PV centroids and cortical pixels were projected onto the nearest point along the pial surface and assigned an arc-length coordinate. The pial surface was subdivided into fixed-width bins, and PV density was computed as cells per mm² by normalizing PV counts to bin area. Median WFA fluorescence was computed across pixels within each bin.

To assess PV-centered PNN localization (Fig. 4G), WFA enrichment was calculated for each PV cell as the difference between median WFA intensity in a perisomatic annulus (5–15 µm from the centroid) and median intensity in a surrounding local background region. A depth-matched null distribution was generated by sampling random cortical points matched to the depth distribution of PV cells and computing the same metric.

## Data and Materials Availability

Code is available on GitHub (https://github.com/ryan-gorzek/opossum-V1-omics). Raw and processed snRNA-seq (GSE299387) and Stereo-seq (GSE299386) data are available on NCBI Gene Expression Omnibus. Immunofluorescence images will be deposited onto Figshare.

## Acknowledgments

We thank S. Jain, F. Xie, D.L. Ringach, and S.L. Zipursky for analytical support and critical feedback. We also thank the UCLA Technology Center for Genomics (TCGB) and Bioinformatics and Translational Pathology Core Laboratory (TPCL), as well as E. Tring, S. Jain, and J. Yoo for experimental support. We used computational and storage services associated with the Hoffman2 Cluster, which is operated by the UCLA Office of Advanced Research Computing’s Research Technology Group. Select images were obtained from BioRender. This work was supported by National Institutes of Health grant R01 EY023871 (JTT).

---

**Figure S1.**
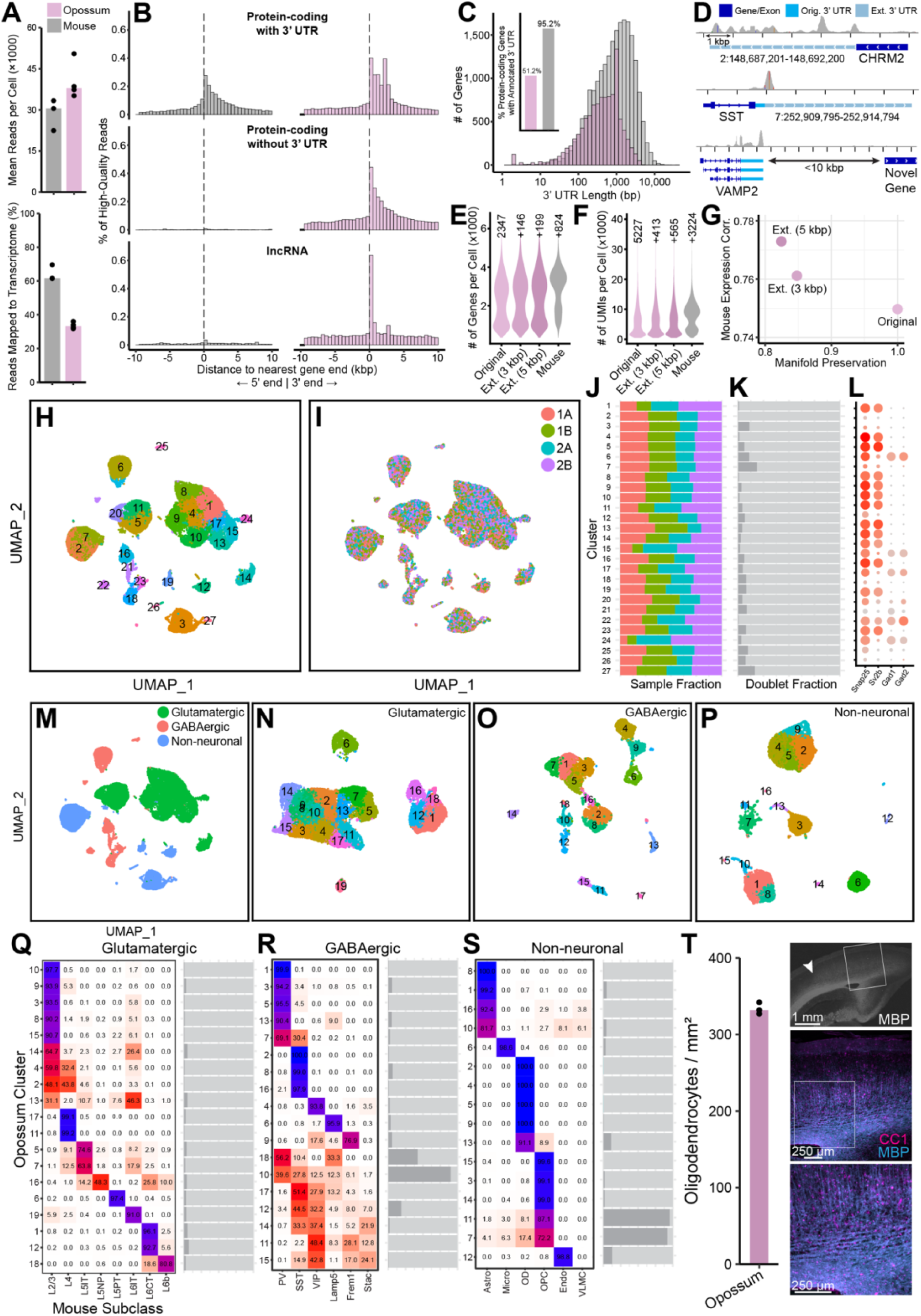
Opossum snRNA-seq data preprocessing. (**A**) Despite comparable sequencing depths, the percentage of snRNA-seq reads mapping confidently to the transcriptome (exons, introns, and untranslated regions [UTRs]) in opossums was nearly half of that in mice. (**B**) Intergenic (mapping confidently to regions outside of exons, introns, or UTRs) read locations relative to the closest gene (negative is 5’ or upstream, positive is 3’ or downstream) in mice and opossums. Distributions are shown by gene biotype (protein-coding with or without an annotated 3’ UTR, or long noncoding RNA [lncRNA]) of the closest gene, as a fraction of total high-quality reads. Because most mouse protein-coding genes have an annotated 3’ UTR (see panel C), most intergenic reads fall into that group. Note the bimodality of that group in opossums (related to panel C). (**C**) Fraction of genes with an annotated 3’ UTR in both species, and distribution of their lengths. Note small spikes at 3 bp (stop codons) and 1000 bp (consistent with annotation artifacts). (**D**) Examples of gene extension that capture downstream read pileups in opossums, even when 3’ UTR annotation is already present (e.g., SST). Genes less than 10 kbp upstream of a neighboring gene were not extended (e.g., VAMP2). (**E–F**) Distribution of genes (E) and transcripts (UMIs; F) per cell in original and extended (3 or 5 kbp) opossum genome configurations, relative to mice. The median is shown for the original opossum genome, with others shown as relative increases. (**G**) Manifold preservation plotted against mouse expression correlation for opossum genome configurations. Manifold preservation was quantified as the mean fraction of each cell’s k=30 nearest neighbors in PCA space (30 PCs) that are shared between the original and extended genomes: a value of 1.0 indicates identical local structure. Mouse expression correlation was computed as the mean Spearman correlation coefficient between opossum and mouse pseudobulk expression profiles aggregated by cell class (glutamatergic, GABAergic, non-neuronal). (**H**) Unsupervised Leiden clustering of all opossum cells passing initial quality control (see Methods) in UMAP space. (**I–J**) UMAP (I) and cluster-wise (J) representations of samples in opossum snRNA-seq data. Numbers (1 and 2) represent biological replicates and letters (A and B) represent technical replicates. (**K**) Cluster-wise representations of putative doublets (see Methods). (**L**) Canonical neuronal (*Snap25*), glutamatergic (*Sv2b*), and GABAergic (*Gad1*, *Gad2*) marker genes used to divide unsupervised clusters into cell classes. (**M**) All opossum cells divided into cell classes. (**N–P**) Unsupervised clustering of glutamatergic (N), GABAergic (O), and non-neuronal (P) cells prior to removal of ambiguous or doublet-enriched clusters. (**Q– S**) Cluster-wise cross-species correspondence (left) and doublet representation (right) for glutamatergic (Q), GABAergic (R), and non-neuronal (S) classes used to guide removal of ambiguous/artifact clusters. (**T**) Oligodendrocyte densities in opossum V1 (left) were measured in sections co-labeled for CC1 (mature oligodendrocytes; magenta) and myelin basic protein (MBP; cyan) marking myelinated processes (right). Related to Fig. 1K, see also (*41*).

**Figure S2.**
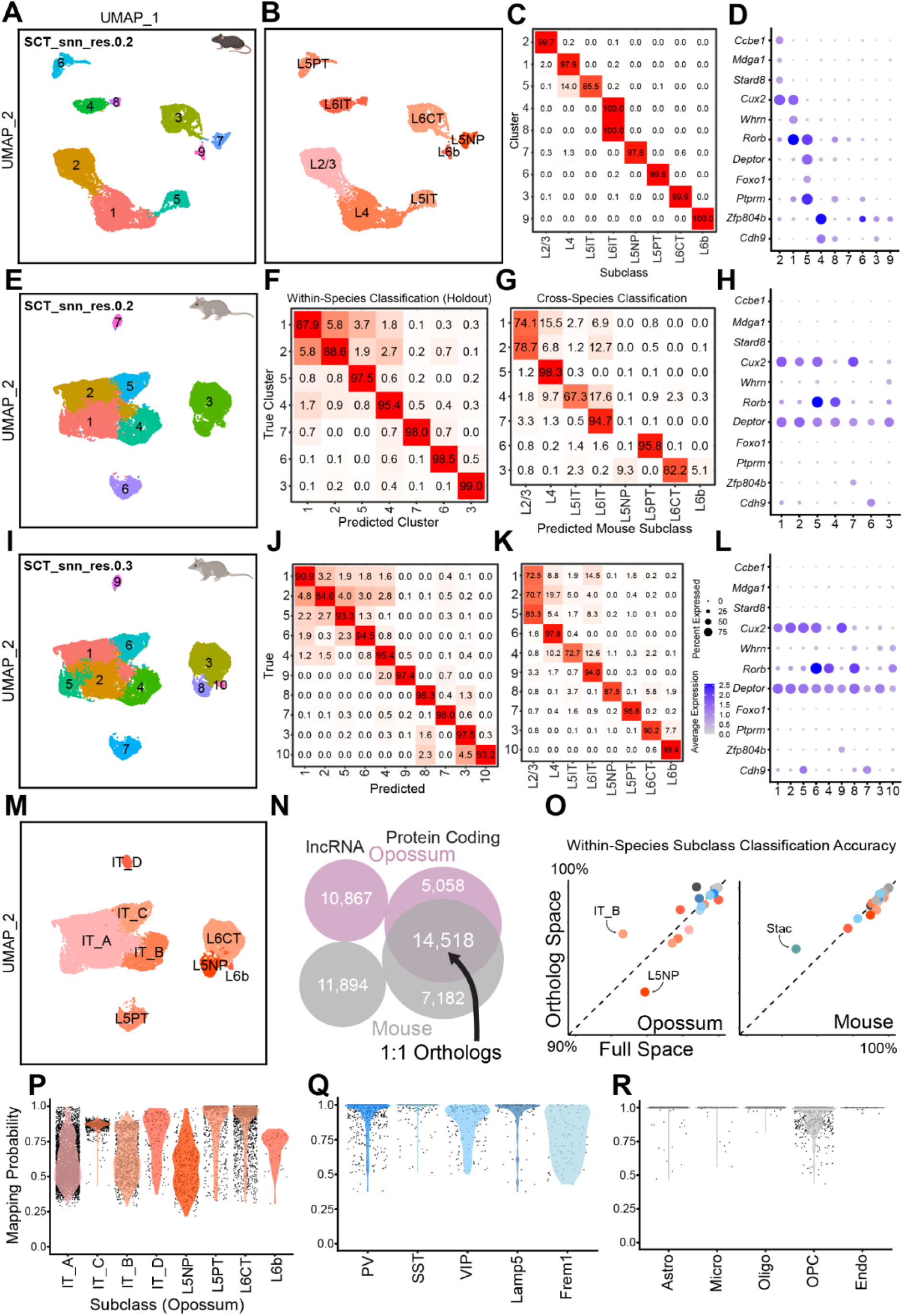
Semi-supervised labeling of opossum intratelencephalic (IT) subclasses and cross-species correspondence. (**A**) Mouse glutamatergic cells in UMAP space, clustered at low resolution (0.2) with the Leiden algorithm. (**B**) Mouse glutamatergic cells in UMAP space, labeled by subclass (*30*). (**C**) Confusion matrix between low-resolution unsupervised clustering and subclass labels in mice. (**D**) Canonical glutamatergic subclass marker gene expression in unsupervised clusters shown in (A). (**E**) Opossum glutamatergic cells in UMAP space, clustered at low resolution (0.2). (**F**) Confusion matrix derived from 50% train-test splits of unsupervised clusters shown in (E). (**G**) Confusion matrix derived from cross-species label transfer (see Methods) between mouse glutamatergic subclasses and unsupervised clusters shown in (E). (**H**) Canonical glutamatergic subclass marker gene expression in unsupervised clusters shown in (E). (**I–L**) Same as (E–H), but with Leiden resolution set to 0.3. (**M**) Final glutamatergic subclass assignment in opossums. Clusters 1 and 2 in (E) were merged to form IT_A. L6CT, L5NP, and L6b correspond to clusters 3, 8, and 10 in (I). (**N**) Composition of the opossum genome. Long noncoding RNAs (lncRNAs) are species-specific and one-to-one orthologs are exclusively protein-coding genes. (**O**) Classification accuracy of 50% train-test splits for each subclass in species-specific (full) and orthologous gene expression space in opossums and mice. (**P–R**) Cross-species mapping quality (obtained from MapQuery, see Methods) scores for opossum cells shown by glutamatergic (P), GABAergic (Q), and non-neuronal (R) subclasses. Related to Fig. 2C–F.

**Figure S3.**
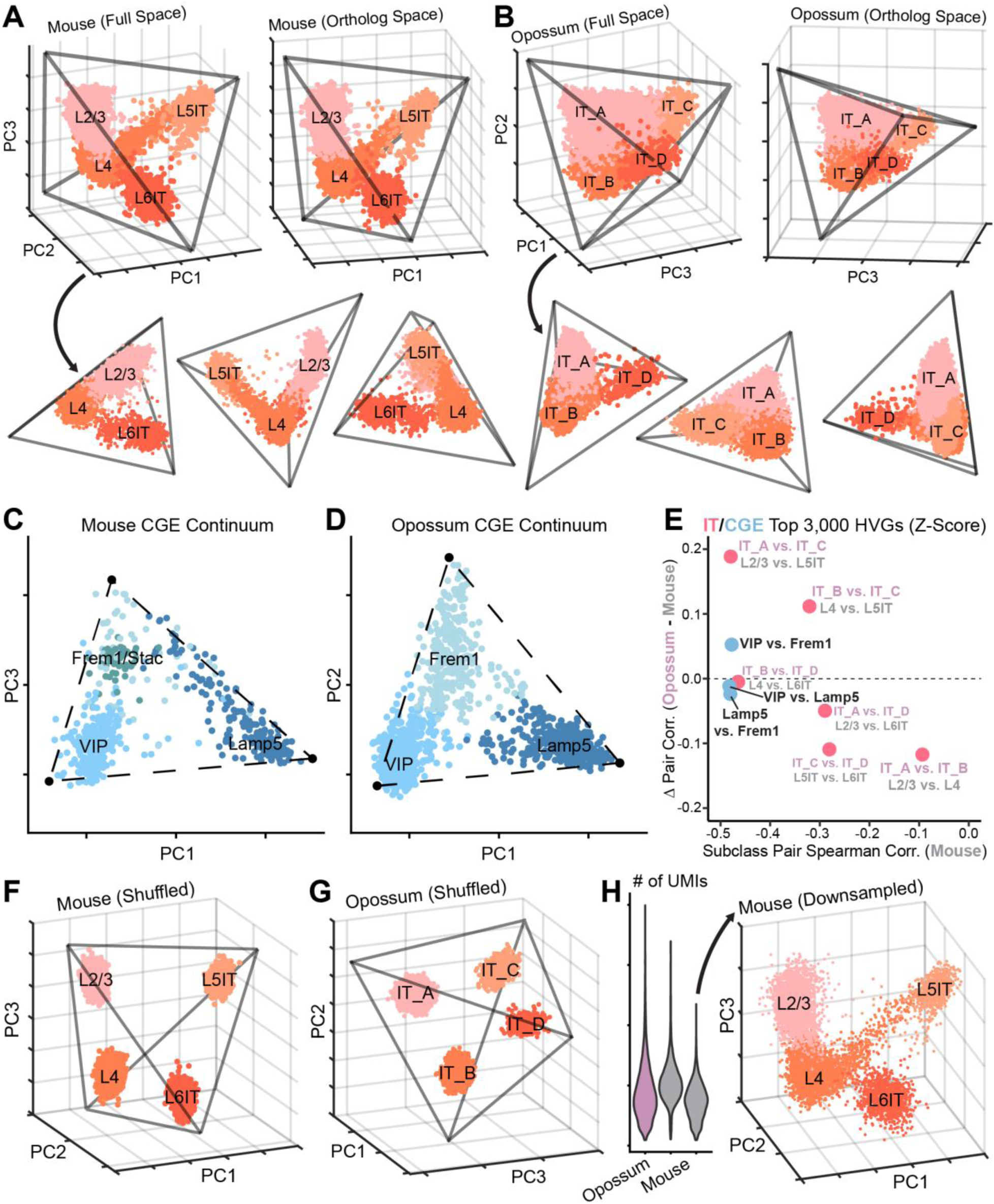
IT principal component (PC) spaces, other gradients, and quality controls. (**A–B**) Tetrahedral IT gradients in mouse (A) and opossum (B) principal component spaces, shown from various angles. Tetrahedrons were fit using established algorithms (see Methods). Related to Fig. 2G–H. (**C–D**) GABAergic neurons derived from the caudal ganglionic eminence (CGE), which form a continuum in gene expression space, display similar triangular gradients in mice (C) and opossums (D). (**E**) Relationship between within-species subclass similarity and cross-species divergence for IT and CGE continua. Each point represents a pair of IT or CGE subclasses. The x-axis shows the Spearman correlation of subclass-pair gene expression in mice, computed over the top 3,000 highly variable genes (z-scored within IT or CGE subclasses). The y-axis shows the difference in this pairwise correlation between opossum and mouse (opossum − mouse). Positive values indicate increased similarity in opossum relative to mouse, while negative values indicate reduced similarity. Note that CGE subclass pairs have similar within-continua similarity to L2/3 and L5IT, but this correlation (IT_A vs. IT_C) increases in opossums considerably more than the CGE pairs. (**F–G**) Randomly shuffling gene expression within subclasses (see Methods) confirms that mouse (F) and opossum (G) IT cells form continua that are not driven by noise (see Methods). (**H**) The mouse IT continuum retains its structure after downsampling UMI (transcript) counts to match opossum levels (see Methods).

**Figure S4.**
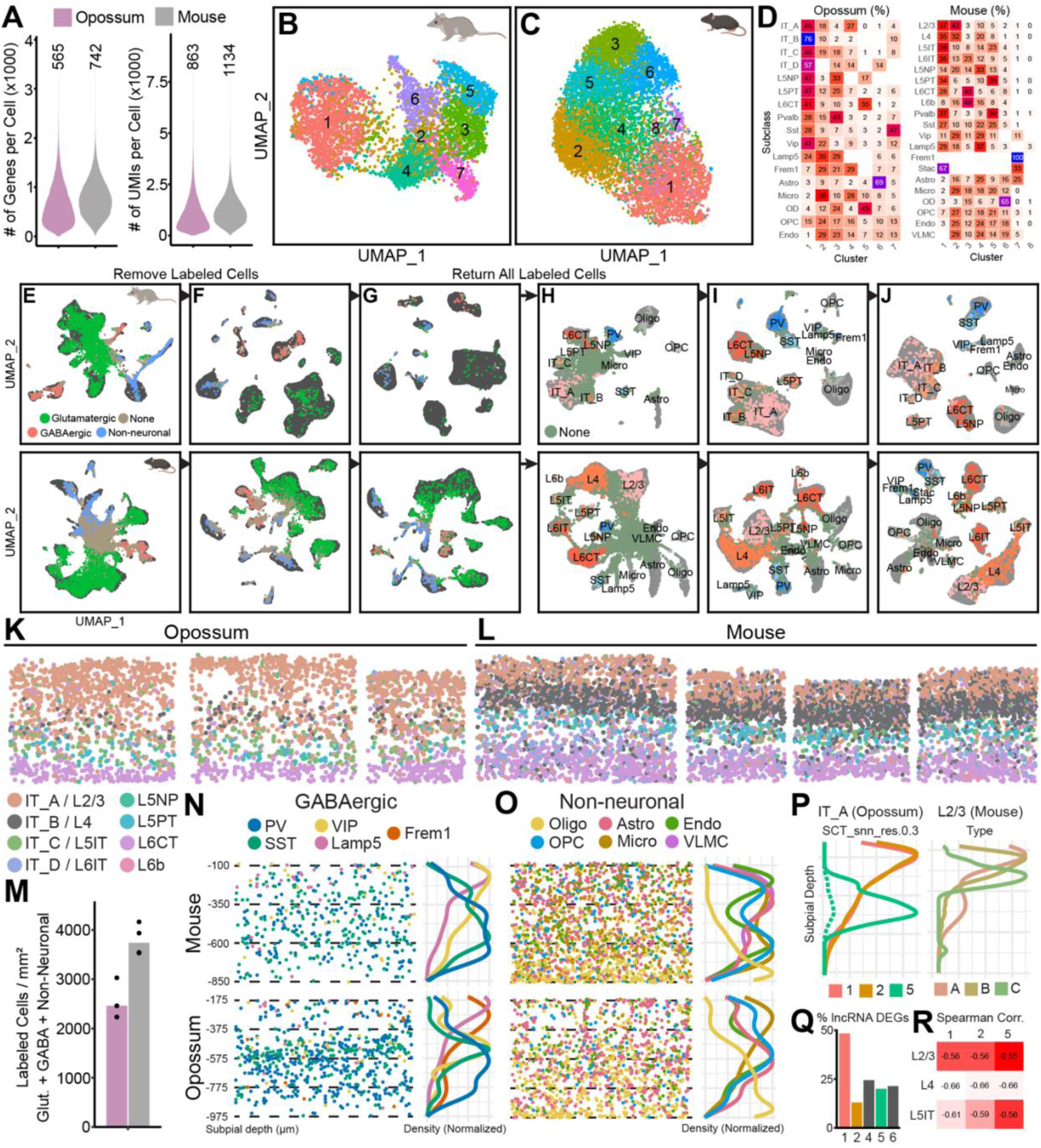
Stereo-seq preprocessing and non-glutamatergic cell subclass distributions. (**A**) Number of genes (left) and UMIs (right) per cell in all Stereo-seq cells from mice and opossums. (**B–C**) Unsupervised clustering of opossum (B) and mouse (C) Stereo-seq cells. (**D**) Final subclass (the result of H–I, see Methods) membership among unsupervised clusters in the original Stereo-seq space (B–C) for opossums (left) and mice (right). (**E–G**) UMAP representation of integrated snRNA-seq (gray) and Stereo-seq (colored by class) cells in opossums (top) and mice (bottom). Stereo-seq class labels were assigned using nearest neighbors in PCA space (see Methods). (**H–J**) Same as (E–G), but with Stereo-seq cells labeled by subclass. Ambiguous cells labeled ‘None’ in (G) were removed before the integration step in (H) (see Methods). (**K–L**) Individual cortical columns extracted from opossum (K) and mouse (L) Stereo-seq sections. Only glutamatergic cells are shown, colored by subclass. (**M**) Total cell density (cells per mm^2^) for mouse and opossum Stereo-seq data following class assignment (see Methods). (**N–O**) Distribution of non-neuronal (N) and GABAergic (O) subclasses from pooled V1 regions. Related to Fig. 3C–D. (**P**) Spatial distributions of IT_A subclusters at Leiden resolution 0.3 (left; fig. S2I–L) and mouse L2/3 types (right). The dashed line for subcluster 5 is normalized to the peak of cluster 1. (**Q**) Fraction of IT subcluster DEGs that are long non-coding RNAs (lncRNAs) at Leiden resolution 0.3 (fig. S2I–L). See Methods for further discussion. (**R**) Spearman correlation between the top 3,000 highly variable genes (HVGs) of opossum IT_A subclusters (Leiden resolution 0.3) and mouse IT subclasses. See Methods for further discussion.

**Table S1.**
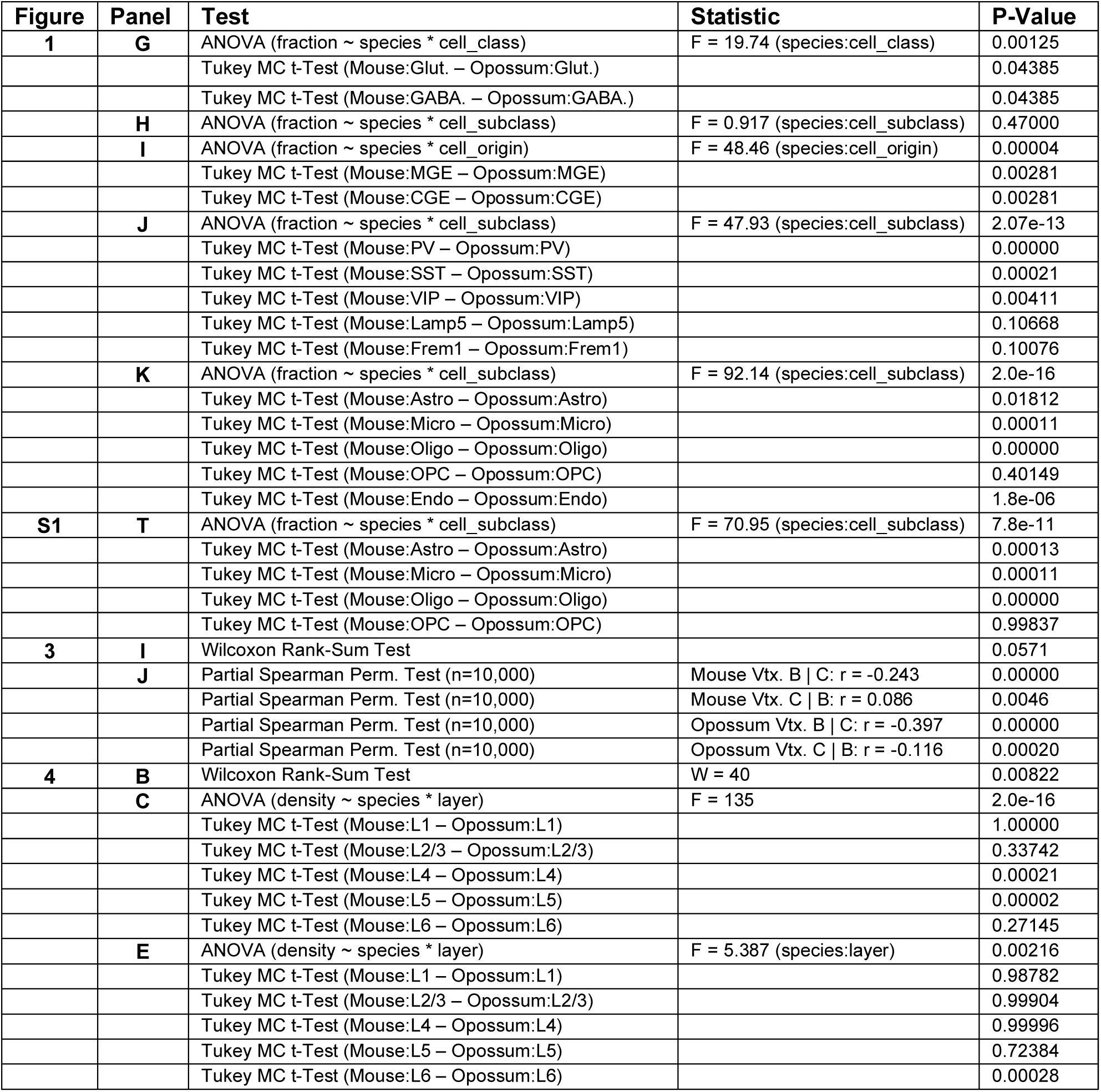
Statistical tests, test statistics, and p-values for all main and supplementary figures.

